# Variation in density, immune gene suppression, and co-infection outcomes among strains of the aphid endosymbiont *Regiella insecticola*

**DOI:** 10.1101/2022.08.28.505589

**Authors:** Elliott B. Goldstein, Yazmin de Anda Acosta, Lee M. Henry, Benjamin J. Parker

## Abstract

Many insects harbor heritable beneficial microbes that influence host phenotypes. Across systems, divergent symbiont strains have been shown to establish at different densities within hosts. This genetic variation is important evolutionarily because within-host density has been linked to the costs and benefits of the symbiosis for both partners. Studying the factors shaping within-host density is therefore important to our broader understanding of host-microbe coevolution. Here we focused on different strains of *Regiella insecticola*, a facultative symbiont of aphids. We first demonstrated that different strains of *Regiella* consistently establish in pea aphids at drastically different densities. We then found that variation in density is correlated with the expression levels of two key insect immune system genes (phenoloxidase and hemocytin), with immune gene suppression correlating with higher *Regiella* density. We then performed an experiment where we established co-infections of a higher- and lower-density *Regiella* strain, and we showed that the higher-density strain is better able to persist in co-infections than the lower-density strain. Together, our results point to a potential mechanism for strain-level variation in symbiont density in this system, and further suggest that symbiont fitness may be increased by establishing at higher density within hosts. Our work highlights the importance of within-host dynamics shaping symbiont evolution.

## Introduction

Most insects harbor heritable bacteria (1). These symbiotic associations can range from parasitic to mutualistic, and symbionts have been shown to influence host traits including host plant use, body color, and defense against natural enemies (2, 3). An important aspect of insect-symbiont interactions is the within-host density of symbiont infections (4-6). Symbiont density has been shown to vary across individual insects, driven in part by genetic variation among microbes and/or hosts (7). For example, the density of *Wolbachia*, a widely-distributed genus of bacteria that often acts as a reproductive parasite, varies drastically across bacterial strains and host species. The molecular mechanisms underlying this variation have been uncovered in some *Wolbachia-*host interactions: a region of the bacterial genome termed Octomom controls density in *Drosophila melanogaster* (8) and a host gene of unknown function drives *Wolbachia* density across species of *Nasonia* wasps (9).

Within-host density is a particularly important aspect of host-symbiont interactions because it has been clearly linked with the fitness of both partners. Higher density infections can confer benefits to hosts in some systems but also impose stronger costs on hosts. For example, higher density *Wolbachia* infections confer stronger protection to *Drosophila* hosts against viral infection (10) but also decrease host survival (11). For symbionts, density has also been shown to influence fitness by impacting transmission success—the density of *Wolbachia* in naturally infected *Drosophila innubila* flies, for example, is correlated with maternal transmission rates. Density might also influence microbial fitness to the extent that the fitnesses of both partners are linked by vertical transmission. However, it is possible that the ‘optimal’ density might be different from the perspectives of host and symbiont fitness, leading to antagonism over control of symbiont density even in mutualistic symbioses. Studying the forces shaping symbiont density, and in particular how symbionts balance within-host selective pressures vs. the effects they have on host fitness, is important for our broader understanding of host-microbe interactions.

Pea aphids (*Acyrthosiphon pisum*) are an important model for studying host-symbiont biology. Aphids host several species of gram-negative bacteria, including an obligate symbiont called *Buchnera aphidicola* that is required for host survival and several ‘facultative’ symbionts that are found at intermediate frequencies in natural populations and can confer important benefits to hosts (2). For example, the Gammaproteobacteria *Regiella insecticola* (hereafter *Regiella*) has been shown to make pea aphids more resistant to specialist fungal pathogens (12, 13). *Regiella* has also been linked to host plant use (14) (though other studies have failed to find these same effects (15-17)), and a strain of *Regiella* isolated from *Myzus persicae* (referred to in this study as strain .515) was shown to play a role in wasp resistance (18). *Regiella* found in natural populations of pea aphids form two main phylogenetic groups (termed “clade 1” and “clade 2”) that are separated by around a half-million years (19). Recent studies have demonstrated extensive horizontal transmission of *Regiella* across host lineages (19), and have even found that individual aphids can harbor co-infections of ‘clade 1’ and ‘clade 2’ *Regiella* strains (20). In previous work, we have found that *Regiella* strains vary in the density at which they establish in the same host genetic background (21). We have also found that *Regiella* density is correlated with the survival costs inflicted on aphids, but not with the protective benefits to aphids from harboring symbionts (21).

Several recent studies have found that some aphid facultative symbionts interact with the innate immune system (22). Specifically, aphids harboring *Regiella* have decreased numbers of circulating immune cells (23) and reduced expression (24) and protein levels of phenolxoidase (25), a critical component of the insect innate immune system. Suppression of phenoloxidase (PO) using RNAi leads to an increase in *Regiella* densities (24). Similar trends have been found for other aphid endosymbionts: a recent study identified specific strains of the aphid facultative symbiont *Hamiltonella defensa* that reach extremely high densities within hosts and suppress host immune genes (22). Together these studies suggest differences in how symbionts interact with the host immune system could contribute to variation in density across symbiont strains.

In this study, we established multiple strains of *Regiella* from pea aphids and two *Regiella* strains from other aphid species (*Myzus persicae* and *Diuraphis noxia*) in a single genotype of pea aphid. We show that variation in *Regiella* density is negatively correlated with the expression of two immune genes across symbiont strains. In particular, we found that ‘Clade 2’ *Regiella* strains establish at high densities in hosts and suppress host immunity, even though in previous work we have found that both clades can provide strong protection against fungal pathogens (26). We then explored why high density *Regiella* strains might persist in natural populations. We established aphid lines with co-infections of a higher-density clade 2 strain and a lower-density clade 1 strain, and we tracked symbiont loss over 8 generations using a strain-specific PCR assay. Our results suggest that the higher-density strain is maintained better in co-infections than the lower-density strain. Together, we argue that suppressing host immunity to establish at high densities might benefit symbionts even at the expense of host fitness, and we discuss these results in light of the within-host vs. host-level selective pressures that shape symbiont evolution.

## Materials and Methods

### Aphid maintenance

We maintained stock lines of aphids on fava bean plants (*Vicia faba* var Windsor) at low densities (<7 adults per plant) at 20°C, 16L:8D. These light and temperature conditions ensure parthenogenetic reproduction of aphids, allowing us to maintain different aphid genotypes in the lab. All experiments in this study used the LSR1-01 aphid line, which was collected from alfalfa (*Medicago sativa*) near Ithaca, NY, US in 1998 and was sequenced in the pea aphid genome project (27). This line was originally collected with *Regiella* (strain .LSR) but was subsequently cured of facultative symbionts using antibiotics.

### Secondary symbiont establishment

We established facultative symbionts into new aphid lines with microinjection using established protocols. Briefly, we attached a glass-pulled needle to a syringe and extracted hemolymph from an adult donor aphid. We then injected hemolymph into one-day-old recipient aphids (1^st^ instars) in the thorax. When the injected aphids became adults, we discarded the first 15 offspring, reared late birth-order offspring to adulthood, and screened them for symbionts. We re-screened each line for *Regiella* before use in experiments where applicable.

### Screening for and genotyping *Regiella*

To detect the presence of *Regiella*, we extracted DNA from whole adult aphids using ‘Bender’ buffer with ethanol precipitation and wash as in previous studies (28). We then used PCR with published *Regiella-*specific primers (19) (94°C 2 min, 11 cycles of 94°C 20s, 56°C (declining 1°C each cycle) 50 s, 72°C 30 s, 25 cycles of 94°C 2 min, 45°C 50 s, 72°C 2 min and a final extension of 72°C 5 min). To assign *Regiella* strains to phylogenetic clades, we amplified and sequenced the *murE* gene as in previous studies (19, 26).

### *Regiella* density in *vivo* using qPCR

We used quantitative PCR (qPCR) to measure *Regiella* within-host density. We used primers that amplify a conserved region of the single-copy *hrpA* gene (F:CGCATTGGGAGAAAAGCCAAG; R:CCTTCCACCAAGCCATGACG) (21, 24). For each sample, we also amplified glyceraldehyde-3-phosphatedehydrogenase (*g3PDH;* ACYPI009769) in the aphid genome, as in previous work (24). For this study, we generated qPCR standards for both genes by cloning the target amplicons into One Shot TOP10 competent *E. coli* cells using the TOPO TA cloning kit with pCR 2.1 vector and extracting amplified plasmids using the GE Healthcare illustra plasmidPrep Mini Spin Kit under recommended conditions. The cloned fragments were sequenced with M13F primer for confirmation and quantified using a nanodrop. We then included a 1:5 serial dilution ranging from 1.6 × 10^7^ to 5.12 × 10^3^ copies of the plasmid on each plate, which allowed us to measure the absolute number of copies of *hrpA* and *g3PDH* in each reaction. We then calculated *Regiella* density by taking the ratio of *Regiella* to host cells using the ratio of *hrpA* to *g3PDH* copies. To analyze variation in *Regiella* density across genotypes, we performed a non-parametric ANOVA (Kruskal-Wallis) in R v.4.0.2 using the kruskal.test function (relative density deviated significantly from a normal distribution). We performed a post-hoc comparison of genotypes using a Dunn test in the FSA package (29).

### Host immune gene expression

We extracted RNA by first removing embryos from pooled groups of four adult aphids from four biological replicates of aphids infected with each symbiont genotype. We used Trizol and chloroform with an isopropanol precipitation and ethanol wash to isolate RNA, followed by the Zymo RNA Clean & Concentrator kit with gDNA wipeout with DNase I. We then used the iScript cDNA synthesis kit from Zymo to convert RNA to cDNA under recommended conditions.

We used primers that amplified 80-120bp of two target immune genes of interest (*PO1*: ACYPI004484 and hemocytin: ACYPI003478) designed in a previous study (24). For endogenous controls, we used two genes (*Glyceraldehyde 3-phosphate dehydrogenase (g3PDH): ACYPI009769* and *rpl32: ACYPI000074*). Concentrations of each primer pair were optimized against a 1:10 serial dilution of gDNA (200ng–0.2ng gDNA per reaction) to an efficiency of 100 +/- 10% (*PO1*: 100nM; hemocytin: 100nM; *g3PDH*: 400/350nM F/R; and *rpl32*: 200nM). Reactions were run on a CFX96 Real-Time System machine (BioRad), with an initial step of 95°C for 3 minutes and 40 cycles of 95°C for 10s and 60°C for 30s. We ran three 20μL technical replicates, each of which included a 1X PCR buffer, Mg+2 at 2mM, dNTPs at 0.2mM, EvaGreen at 1X, 0.025 units/μL of Invitrogen taq, and 40ng of cDNA. We analyzed the effects of symbiont genotype on the expression levels of each gene using linear models in R v.4.0.2, with genotype as a fixed effect. We then compared expression in each aphid line harboring *Regiella* with symbiont-free aphids using post-hoc comparisons with the ‘multcomp’ package (30).

To measure correlations between immune gene expression and symbiont density, we calculated the average -ΔΔC_T_ value for PO and for hemocytin for each genotype of *Regiella* (the amount each gene was downregulated compared to symbiont-free aphids), and we compared expression to symbiont density using Spearman’s rank correlation tests, implemented in R v.4.0.2 with the cor.test (method=‘pearson’).

### Strain specific *Regiella* detection

We designed primers that targeted strain-level variation in the *hrpA* gene among *Regiella* genotypes. Each primer pair amplified only one of two strains (genotype .LSR (F:*5’-*GCCCGTTTTGCTGTTTTCCC*-3’* R:5’-AAAAGCATGGCTGGTTTGCC-3’) or genotype .313 (F:5’-GCCCGTTTCGCTATTTTCCC-3’ R:AAAAACGTGACGGGTTTGCC-3’), with PCR conditions of 95°C 2 min, 30 cycles of 95°C 30 s, 65°C 30 s, 68°C 50 s, and a final extension of 68°C 5 min. We verified that these primers were strain specific and could detect *Regiella* co-infections at up to a 1:100 dilution of DNA mixed from the two strains (Figure S1).

### Co-infection assay

We created co-infections of aphids with two strains of *Regiella* (.313 and .LSR). We performed microinjections as above from an adult donor aphid to a 1-2 day old aphid with an existing infection of the other genotype of *Regiella* (.LSR injected into aphids already harboring .313, and vice versa) to create a co-infection with both genotypes. We injected the same volume of hemolymph from donor aphids into each recipient, which means that the number of bacteria injected into recipients differed between .LSR and .313 injections. We therefore also included injections of each strain of *Regiella* into control (uninfected) aphids in the experiment. At each generation after injection (the 1^st^ through 8^th^ generations), we collected a single adult aphid from the cage and tested it for both strains of *Regiella* using the genotype-specific primers described above. Each aphid was determined to have an infection of *Regiella* genotype .LSR, *Regiella* genotype .313, a co-infection with both genotypes, or no detectable *Regiella* infection.

We analyzed these data using generalized linear mixed effects models with the lme4 package in R v.4.0.2 (31). We assessed symbiont maintenance across the experiment, with the presence or absence of the injected symbiont strain as the dependent variable (modeled as a binomial outcome as determined by the presence or absence of PCR amplification). We included two factors and their interaction as fixed effects in the model: the strain of the injected symbiont (strain .LSR from ‘clade 1’, or strain .313 from ‘clade 2’) and the background of the recipient aphid (i.e. whether it had a preexisting *Regiella* of the opposite strain, or symbiont-free). As random effects, we included generation and experimental replicate. We compared models by building minimal models first by removing the interaction term, then aphid background, and then symbiont strain, and we determined statistical significance through model comparisons using ANOVA and a chi-squared test.

## Results

### Symbiont density

We found significant variation in the relative density of *Regiella* strains (Kruskal-Wallis ξ^2^ = 28.5, df = 6, p < 0.0001), with the ratio of *Regiella* / host cells in each aphid ranging from 0.3 to 14.8 (Figure 1A). Specifically we found that the three ‘clade 2’ *Regiella* strains established at higher densities than strain .515, and strain .313 established at a significantly higher density than strain .Md10 from ‘clade 1’ (post-hoc tests at p < 0.05; Figure 1A).

**Figure 1:**
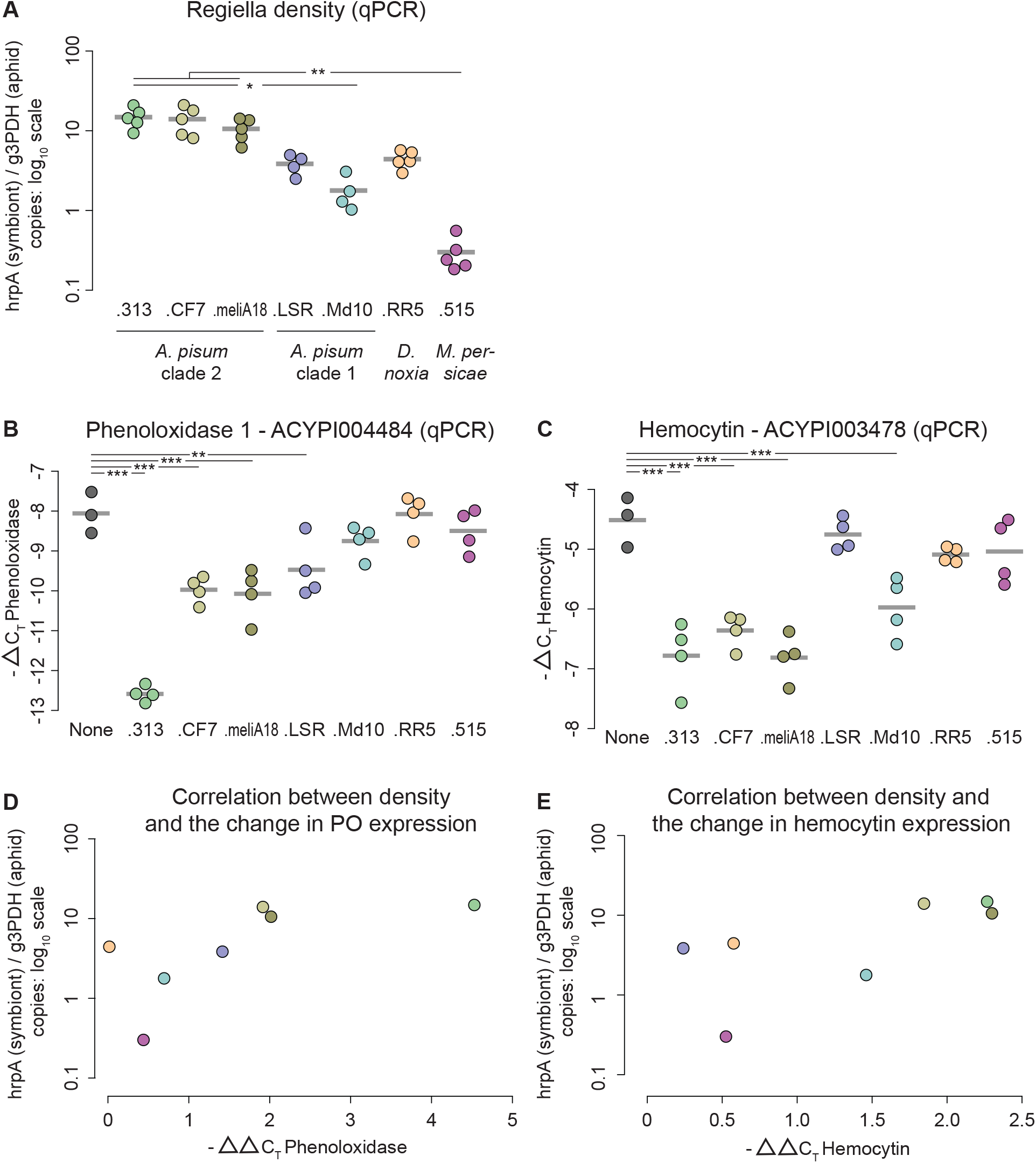
Within-host density and effects on aphid immune gene expression of different *Regiella* strains. **A:** Relative *Regiella* density measured by qPCR. *Regiella* strains are shown along the bottom of the figure, each using a different color. The y-axis shows relative symbiont density as the ratio of *hrpA* copies to *g3PDH* copies. Each sample is shown with a colored point, with the mean of each genotype indicated with a grey bar. Statistical significance using a post-hoc rank-order comparison is shown along the top of the figure. **B&C:** Immune gene expression of aphids harboring each line. Panels B and C show expression of PO1 and Hemocytin, respectively. The y-axes show -ΔC_T_ values of expression (mean endogenous control C_T_ values – target gene C_T_ values). Statistical significance is shown along the top of the figure. **D&E:** Correlation between symbiont density and immune gene expression for PO1 (panel D) and hemocytin (panel E). Density is shown along the y-axes as in A. The x-axes show the average -ΔΔC_T_ values, comparing expression of aphids harboring each symbiont strain vs. uninfected aphids. Values further to the right on the y-axis represent stronger decreases in immune gene expression due to symbiont presence.

### Immune gene expression

Symbiont strain had a significant effect on Phenoloxidase 1 (ACYPI004484) expression (F = 33.9, df = 7, p < 0.0001; Figure 1B), with genotypes .313, .CF7, .meliA18, and .LSR significantly decreasing expression relative to symbiont-free aphids (Table S2, Figure 1B). Similarly, *Regiella* influenced Hemocytin (ACYPI003478) expression (F = 19.1, df = 7, p < 0.0001; Figure 1C), with .313, .CF7, .meliA18, and .MD10 significantly decreasing hemocytin expression (Table S3, Figure 1C).

*Regiella* density was also negatively correlated with expression of Phenoloxidase 1 (t = 3.18, df = 5, p = 0.025; Figure 1D) and with expression of hemocytin (t = 2.92, df = 5, p = 0.033; Figure 1E). Higher density *Regiella* strains were associated with larger decreases in PO1 and hemocytin expression (relative to uninfected aphids).

### Coinjection experiments

We found that symbiont genotype did not influence establishment success (Chisq = 2.1, 1DF, p = 0.15), but whether a host already contained a *Regiella* strain had a strong impact on whether injected symbionts could establish in a host (Chisq = 37.7, 1DF, p < 0.0001). Importantly, there was a significant interaction between *Regiella* genotype and whether recipients already harbored a symbiont (Chisq = 9.3, 1DF, p = 0.0023). Most replicates of strain .LSR *Regiella* were unable to establish in aphids already harboring strain .313 *Regiella*, with lines losing strain .LSR at generation 2 after injection (Figure 2, top). Strain .313 *Regiella*, in contrast, were largely able to establish in aphids that already harbored a strain .LSR infection and persisted throughout the experiment in a co-infection (Figure 2, bottom).

**Figure 2:**
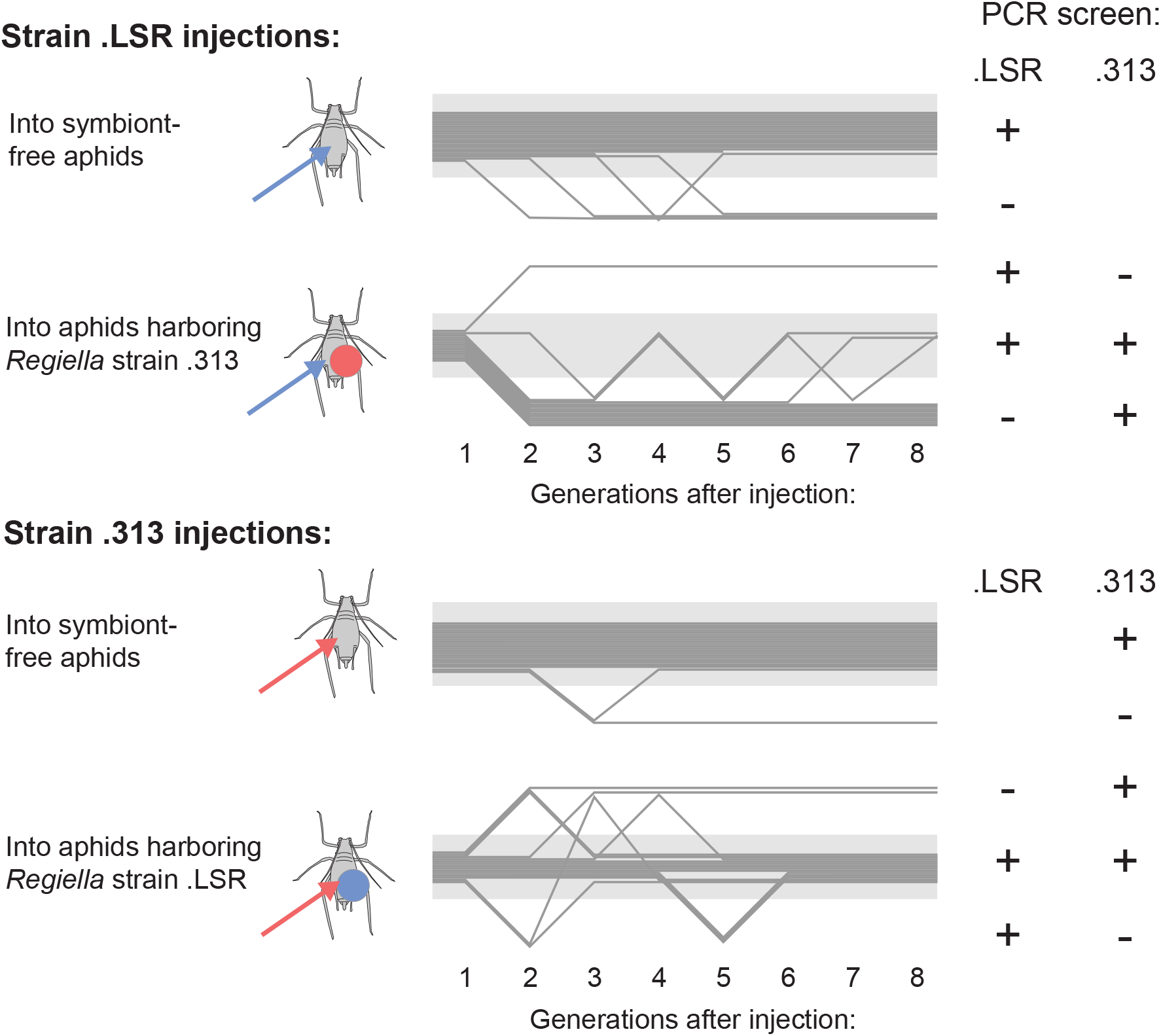
Co-infection outcomes of *Regiella* strains. Each dark grey line in the figures represents a separate aphid line, tracked for 8 generations shown along the x-axis. Injections with strain .LSR (*A. pisum* ‘clade 1’) are shown in the top panel, and injections with strain .313 (*A. piusm* ‘clade 2’) are shown in the bottom panel. Within each panel, recipient aphids either harbored no secondary symbionts (symbiont-free aphids) or harbored *Regiella*. Vertical movement on the plots represents the outcome of PCR screens for *Regiella*. Injections into symbiont-free aphids were either retained or lost at each generation. Co-infections with two *Regiella* strains were either maintained as co-infections, or one of the symbiont strains was lost, as indicated to the right of the figure. Note that at each generation, a single aphid from each host plant was harvested at random for DNA extraction and PCR screening—lines therefore can ‘regain’ a symbiont across generations due to individual differences among aphids on a plant.

## Discussion

In this study, we measured within-host density of multiple *Regiella* strains and the effects of each strain on the expression levels of two host immune genes. We found significant variation in density among *Regiella* genotypes (Figure 1A). In particular, a sequenced strain (32) of *Regiella* that was isolated from another aphid species (*Myzus persicae* strain .515) (18) is harbored at extremely low densities (more than an order of magnitude below other strains) but has been nonetheless transmitted vertically over many years of laboratory propagation.

The aphid immune system has been implicated in the regulation of vertically-transmitted symbioses in several studies, with two related symbionts (*Regiella insecticola* and *Hamiltonella defensa*) leading to decreased expression of key insect innate immune system genes and fewer circulating immune cells (22-25). We have also found that knocking down expression of Phenoloxidase using RNAi leads to increased *Regiella* density over aphid development (24). In this study, we found that strains of *Regiella* vary in their suppressive effects on host immune genes (Figures 1B and 1C), and that stronger suppression is correlated with higher symbiont densities across *Regiella* strains (Figures 1D and 1E). Together these studies point to a potential mechanism for strain-level variation in *Regiella* density, namely that strains might reach higher densities within hosts by more strongly suppressing host immune mechanisms.

We then designed an experiment with the goal of assessing whether co-infecting strains varied in their ability to persist inside hosts across generations. We found that a higher-density strain (.313) and a lower-density strain (.LSR) persisted equally well in single infections across eight aphid generations, but that the higher-density strain was better at persisting in a co-infection of both strains (Figure 2).

Specifically, when the lower-density strain .LSR was injected into aphids already harboring strain .313, the injected symbiont was quickly lost from aphid lines. In contrast, the higher-density strain .313 was able to establish and persist as a co-infection in aphids already harboring strain .LSR.

Harboring *Regiella* can be costly for hosts, and higher density symbiont infections (due to both host and symbiont genotype) impose stronger survival costs on aphids (21). One possibility is that some *Regiella* strains have evolved to reach higher densities within hosts in order to provide stronger protective benefits to aphids against natural enemies. However, we have found that protection against fungal pathogens is highly genotype by genotype dependent in this system (a pattern which also extends to other aphid symbionts interacting with parasitoid wasps (33, 34)), and we have not found that symbiont density and fungal protection are correlated across host or symbiont genetic variation. Further, our data shows that symbiont strains that establish at low densities can be faithfully transmitted over multiple years in the laboratory, and we did not find significant differences in symbiont maintenance between the high and low density *Regiella* strain in our experiment. The evidence therefore suggests that higher density symbiont infections are not beneficial for host aphids.

We suggest that within-host dynamics could contribute to the evolution of high microbial densities in heritable symbioses. One possibility is that strains that reach higher densities in hosts have a competitive advantage over lower-density strains. Another possibility is that higher density strains could have better transmission success overwinter or a higher likelihood of horizontal transfer. Although horizontal transmission of aphid symbionts has been demonstrated on evolutionary (35-38) and even ecological (39) timescales, the mechanism of these transfers is unknown. Possibilities include transmission through parasitism events (39) or through aphid ‘honeydew’ from the surface of host plants.

Our results fit well with other recent studies on symbiont density that have emphasized the importance of within-host dynamics on symbiont evolution. A recent experimental evolution study of *Wolbachia* infections in *Drosophila*, for example, did not find any reduction of symbiont density or in the fitness costs of symbionts to hosts across 17 generations of lab conditions expected to select for reduced symbiont virulence (40). The authors speculate that within-host pressures select for symbionts with higher density despite the increased cost to hosts (5, 40, 41). Together these studies suggest that vertically-transmitted symbionts are subject to the same tradeoffs between virulence and transmission that shape pathogen coevolutionary dynamics, and stress the critical importance of within-host dynamics in the evolution of vertically-transmitted symbioses.

## Supporting information

raw_data

## Data accessibility

All data is included as a supplemental file.

## Acknowledgements

Keertana Tallapragada and Will Brewer provided valuable assistance with the aphid rearing. This work was funded by startup funds from the University of Tennessee and US National Science Foundation (NSF) Grant IOS-2152954 to BJP, and by BBSRC grant BB/W001632/1 to LMH. BJP is a Pew Scholar in the Biomedical Sciences, funded by the Pew Charitable Trusts.

## Author Contributions

EBG, LMH, and BJP conceived of the project. EBG and BJP wrote the manuscript. EBG and YdAA carried out the experiments and molecular work. LMH and BJP contributed reagents, materials, and funding for the project. All authors edited the manuscript and approved the final version for submission.

**Table S1:** Post hoc comparisons of symbiont density.

| Comparison | Z | Adjusted p-value |
| --- | --- | --- |
| 313 - 515 | 4.06 | 0.0011 ** |
| 313 - CF7 | 0.20 | >0.99 |
| 515 - CF7 | -3.9 | 0.0024 ** |
| 313 - LSR | 2.3 | 0.4727 |
| 515 - LSR | -1.5 | >0.99 |
| CF7 - LSR | 2.1 | 0.7565 |
| 313 - Md10 | 3.1 | 0.0477 * |
| 515 - Md10 | -0.77 | >0.99 |
| CF7 - Md10 | 2.9 | 0.0869 |
| LSR - Md10 | 0.73 | >0.99 |
| 313 - MeliA18 | 0.69 | >0.99 |
| 515 - MeliA18 | -3.4 | 0.0159 * |
| CF7 - MeliA18 | 0.49 | >0.99 |
| LSR - MeliA18 | -1.6 | >0.99 |
| 1Md10 - MeliA18 | -2.4 | 0.3396 |
| 313 - RR5 | 2.2 | 0.5973 |
| 515 - RR5 | -1.9 | >0.99 |
| CF7 - RR5 | 2.0 | 0.9671 |
| LSR - RR5 | -0.22 | >0.99 |
| Md10 - RR5 | -1.0 | >0.99 |
| MeliA18 - RR5 | 1.5 | >0.99 |

**Table S2:** Post-hoc analysis of Phenoloxidase expression.

| *Regiella* strain | t value | p-value |
| --- | --- | --- |
| .313 | -11.7 | < 0.001 *** |
| .CF7 | -4.9 | < 0.001 *** |
| .meliA18 | -5.2 | < 0.001 *** |
| .LSR | -3.7 | 0.007 ** |
| .Md10 | -1.8 | 0.32 |
| .RR5 | -0.05 | >0.99 |
| .515 | -1.14 | 0.74 |

**Table S3:**
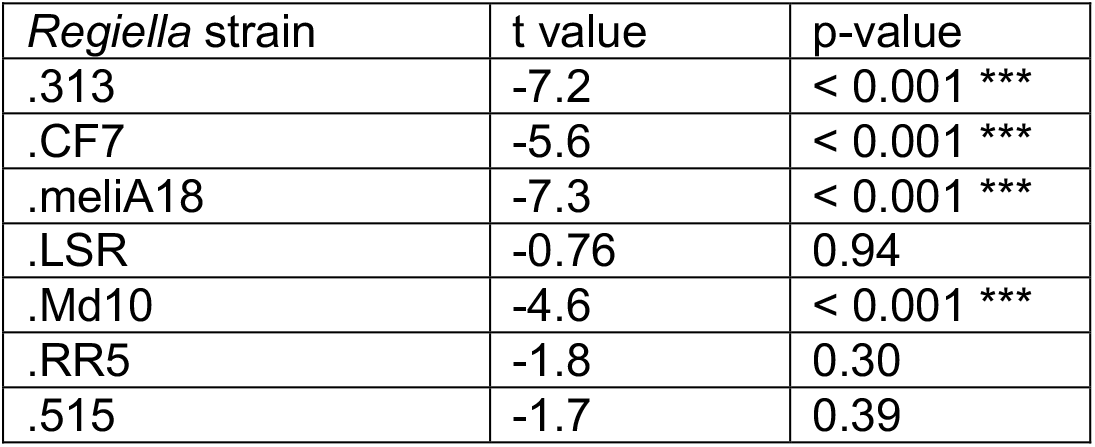
Post-hoc analysis of Hemocytin expression.

| *Regiella* strain | t value | p-value |
| --- | --- | --- |
| .313 | -7.2 | < 0.001 *** |
| .CF7 | -5.6 | < 0.001 *** |
| .meliA18 | -7.3 | < 0.001 *** |
| .LSR | -0.76 | 0.94 |
| .Md10 | -4.6 | < 0.001 *** |
| .RR5 | -1.8 | 0.30 |
| .515 | -1.7 | 0.39 |

**Figure S1:**
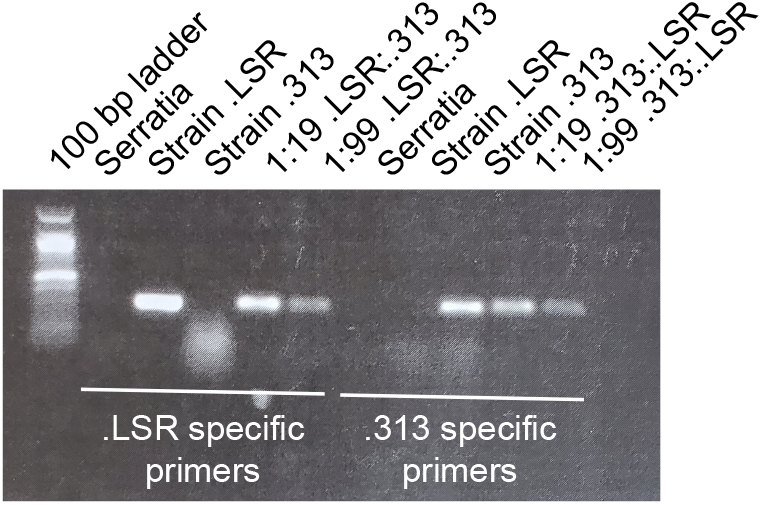
Strain-specific PCR assay. This image shows PCR products run on an agarose gel, with a 100bp ladder to the left of the figure. The first 5 wells are the .LSR specific primers tested against an unrelated symbiont (*Serratia symbiotica*), against DNA from strain .LSR, from .313, and two dilutions of .LSR:.313 DNA at 1:19 and 1:99, respectively. The last 5 wells are the .313 specific primers tested against DNA from *S. symbiotica*, strain .LSR, strain .313, and 1:19 and 1:99 dilutions of .313:.LSR DNA.

